# Evidence for increased fitness of a plant pathogen conferred by epigenetic variation

**DOI:** 10.1101/2023.08.16.553519

**Authors:** Rekha Gopalan-Nair, Aurore Coissac, Ludovic Legrand, Céline Lopez-Roques, Yann Pécrix, Céline Vandecasteele, Olivier Bouchez, Xavier Barlet, Anne Lanois, Alain Givaudan, Julien Brillard, Stéphane Genin, Alice Guidot

## Abstract

Adaptation is usually explained by adaptive genetic mutations that are transmitted from parents to offspring and become fixed in the adapted population. However, more and more studies show that genetic mutation analysis alone is not sufficient to fully explain the processes of adaptive evolution and report the existence of non-genetic (or epigenetic) inheritance and its significant role in the generation of adapted phenotypes. In the present work, we tested the hypothesis of the role of DNA methylation, a form of epigenetic modification, in adaptation of the plant pathogen *Ralstonia solanacearum* to the host plant during an experimental evolution. Using SMRT-seq technology, we analyzed the methylomes of 31 experimentally evolved clones that were obtained after serial passages on a given host plant during 300 generations, either on susceptible or tolerant hosts. Comparison with the methylome of the ancestral clone revealed between 12 and 21 differential methylated sites (DMSs) at the GTWWAC motif in the evolved clones. Gene expression analysis of the 39 genes targeted by these DMSs revealed limited correlation between differential methylation and differential gene expression. Only one gene showed a correlation, the RSp0338 gene encoding the EpsR regulator protein. The MSRE-qPCR (Methylation Sensitive Restriction Enzyme - qPCR) technology was used as an alternative approach to assess the methylation state of the DMSs found by SMRT-seq between the ancestral and evolved clones. This approach also found the two DMSs upstream of RSp0338. Using site-directed mutagenesis, we demonstrated the contribution of these two DMSs in host adaptation. As these DMSs appeared very quickly in the experimental evolution, we hypothesize that such fast epigenetic changes can allow rapid adaptation to the plant stem environment. To our knowledge, this is the first study showing a link between epigenetic variation and evolutionary adaptation to new environment.

## Introduction

Faced with the selection pressure imposed by their environment, pathogens must continuously adapt to survive and multiply. Many works aim to better understand the adaptive processes of pathogens in order to better apprehend the sustainability of the control strategies. Adaptation, the modification of the phenotype as a result of natural selection, is usually explained by adaptive genetic mutations that are transmitted from parents to offspring and become fixed in the adapted population (Lenski 2017; Xue et al. 2019; Gatt and Margalit 2021). However, more and more studies show that genetic mutation analysis alone is not sufficient to fully explain the processes of adaptive evolution and report the existence of non-genetic (or epigenetic) inheritance and its significant role in the generation of adapted phenotypes (Lind and Spagopoulou 2018; Danchin et al. 2019). Epigenetic changes were described to be more involved in short-term adaptation, or acclimation, by inducing phenotypic plasticity (Vogt 2023). This was supported by the observation that epigenetic changes occur at a faster rate than genetic mutations but may be less stable (van der Graaf et al. 2015; Walworth et al. 2021). However, recent works also support the hypothesis that epigenetic modifications could impact long-term adaptive responses to changing environments through the transgenerational inheritance of epigenetic signatures (Kronholm and Collins 2016; Kronholm et al. 2017; Danchin et al. 2019; Stajic et al. 2019; Walworth et al. 2021; Vogt 2023).

A well-documented epigenetic mechanism known to be involved in the modification of the phenotypes is DNA methylation. DNA methylation consists in the addition of a methyl group (CH_3_) on the adenine or cytosine base of DNA catalyzed by DNA methyltransferases (MTases) that will recognize a specific DNA motif. In bacterial genomes, methylated DNA is found in the forms of 6mA (6-methyladenine), which is the most prevalent form; 4mC (4-methylcytosine) and 5mC (5-methylcytosine) (Clark et al. 2012; Blow et al. 2016). Many works demonstrated the role of DNA methylation in the regulation of important cellular functions in bacteria, including DNA replication, DNA repair, chromosome segregation, transcriptional regulation, phenotypic heterogeneity and virulence (Casadesús and Low 2006; López-Garrido and Casadesús 2010; Estibariz et al. 2019; Nye et al. 2019; Payelleville et al. 2019; Sánchez-Romero and Casadesús 2020; Oliveira and Fang 2021). Nowadays, thanks to the Pacbio sequencing technology enabling sequencing of single molecules in real time (SMRT-seq) without amplification, it is now possible to analyze the global DNA methylation profile (methylome) of bacteria (Clark et al. 2012; Murray et al. 2012; Davis et al. 2013; Blow et al. 2016; Beaulaurier et al. 2019). Here, we used SMRT-seq technology to explore the methylome of the model bacterial plant pathogen *Ralstonia solanacearum.* The purpose of this study was to test the hypothesis of methylome variation during an experimental adaptation of the bacteria to various host plants and the potential role of methylome changes in the generation of adapted phenotypes.

*R. solanacearum* is a soil-born plant pathogen responsible of the lethal bacterial wilt disease on more than 250 plant species including economically important crops such as tomato, potato or banana (Vailleau and Genin 2023). This bacterium is worldwide distributed and represents a major threat in agriculture. It is characterized by a strong adaptive capacity, no effective control method is available today and new strains able to colonize new hosts are continuously emerging (Wicker et al. 2007; Wicker et al. 2009; Lopes et al. 2015; Jiang et al. 2016; Bergsma-Vlami et al. 2018). Many works investigated in a better understanding of the adaptive processes in *R. solanacearum*. The role of genetic modifications of the bacterial genome such as mutation, transposable elements movement, recombination or horizontal gene transfer were reported (Coupat-Goutaland et al. 2011; Wicker et al. 2012; Lefeuvre et al. 2013; Guidot et al. 2014). However, the contribution of epigenetic modifications in *R. solanacearum* adaptation has not yet been addressed.

A recent study compared the methylomes using SMRT-seq of two *R. solanacearum* strains belonging to distant phylogenetics groups, the GMI1000 strain from phylotype I and the UY031 strain from phylotype II (Erill et al. 2017). This work identified a commonly methylated motif in the two strains, the GTWW<u>A</u>C motif, 6mA methylated, associated with a MTase, M.RsoORF1982P, that is conserved in all complete *Ralstonia* spp. genomes and across the *Burkholderiaceae* (Erill et al. 2017). Analysis of the methylated regions in *R. solanacearum* genomes identified genes involved in global and virulence regulatory functions thus suggesting a role of DNA methylation in regulation of their expression.

In our previous works, we conducted an experimental evolution of the *R. solanacearum* GMI1000 strain in order to better understand the molecular bases of adaptation. In this experiment, strain GMI1000 was maintained in a fixed plant line during 300 generations by serial passages from stem to stem. This experiment was conducted on six different plant species including susceptible hosts (tomato var. Marmande, eggplant var. Zebrina, pelargonium var. Maverick Ecarlate) and tolerant hosts (bean var. Blanc Précoce, cabbage var. Bartolo, tomato var. Hawaii 7996) (Guidot et al. 2014; Gopalan-Nair et al. 2020). Most of the evolved clones showed a better fitness in their experimental host than the ancestral clone. Whole genome sequence analysis revealed between zero and three mutations in the adapted clones and the role of some mutations in host adaptation was demonstrated (Guidot et al. 2014; Perrier et al. 2016; Perrier et al. 2019; Gopalan-Nair et al. 2020). However, in several adapted clones no mutation could be detected, suggesting that epigenetic modifications may play a role in host adaptation. In addition, transcriptomic analysis of these clones revealed important differential gene expression compared to the ancestral clone, thus reinforcing the hypothesis of a role of epigenetic modification in gene expression change (Gopalan-Nair et al. 2020; Gopalan-Nair et al. 2023).

In this study, we analyzed the methylomes of 31 experimentally evolved clones using SMRT-seq. Comparison with the methylome of the ancestral GMI1000 clone revealed differential methylated sites (DMSs) at the GTWWAC motif in the evolved clones. Using site-directed mutagenesis, we demonstrated the contribution of one DMS in host adaptation, which, interestingly, turns out to be linked to a gene involved in the expression of a bacterial virulence determinant.

## Materials and methods

### Bacterial strains, plant material and growth conditions

The GMI1000 strain and the 31 derived evolved clones investigated in this study are described in table 1. The evolved clones generated after experimental evolution include ten clones evolved in tomato Hawaii 7996 (*Solanum lycopersicum*) (Gopalan-Nair et al. 2020), seven clones in eggplant MM61 (*S. melongena* var. Zebrina), three clones in bean (*Phaseolus vulgaris* var. Blanc Précoce), six clones in tomato Marmande (*S. lycopersicum* var. Super Marmande), and five clones in cabbage (*Brassica oleracea* var. Bartolo) (Guidot et al. 2014). The bacterial strains were grown at 28°C (under agitation at 180 rpm for liquid cultures) either in BG complete medium or in MP synthetic medium (Plener et al. 2010). The pH of the MP medium was adjusted to 6.5 with KOH. For agar plates, BG medium was supplemented with D-Glucose (5 g/l) and triphenyltetrazolium chloride (0.05 g/l). The MP medium was supplemented with L-Glutamine (10 mM) and oligo elements (1000 mg/l) (Gopalan-Nair et al. 2023).

**Table 1.**
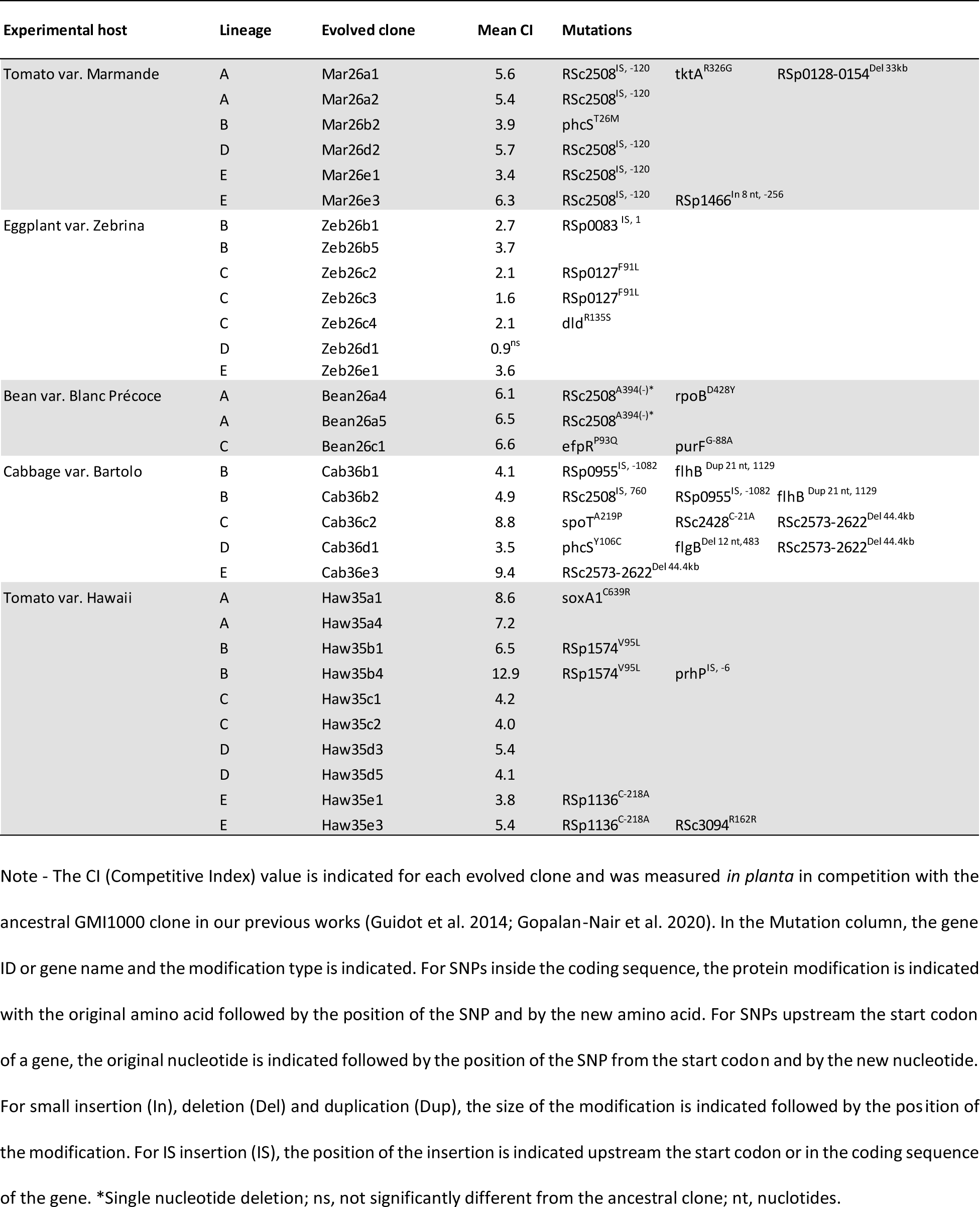
Investigated evolved clones (derived from Gopalan-Nair et al., 2023)

Four to five-week-old tomato (*Solanum lycopersicum*) cultivar Marmande plants were used for the *in planta* bacterial competition assays. Tomato plants were grown in a greenhouse. *In planta* competition experiments were conducted in a growth chamber under the following conditions: 12 h light at 28°C, 12 h darkness at 27°C and 75% humidity.

### SMRT-seq

Genomic DNA was prepared from the bacterial cells grown in synthetic media with glutamine collected at the beginning of stationary phase in order to limit the number of cells in division and avoid a bias towards hemimethylated marks. The bacterial samples were collected as described previously (Gopalan-Nair et al. 2020). Briefly, each of the evolved clones and the ancestral clone GMI1000 were grown in MP medium with 10mM glutamine. For whole genome sequencing, 20 ml of the bacterial culture was centrifuged at 5000g for 10 minutes followed by washing the pellets with water and centrifuged again. The pellets were stored at -80°C until DNA extraction. The DNA were prepared based on the protocol described for high molecular weight genomic DNA (Mayjonade et al. 2017).

Library preparation was performed at GeT-PlaGe core facility, INRAE Toulouse, France and SMRT sequencing at Gentyane core facility, INRAE Clermont-Ferrand, France. Eight libraries of multiplex samples were performed according to the manufacturer’s instructions “Procedure-Checklist-Preparing-Multiplexed-Microbial-SMRTbell-Libraries-for-the-PacBio-Sequel-System.” At each step, DNA was quantified using the Qubit dsDNA HS Assay Kit (Life Technologies) and DNA purity was tested using the nanodrop (Thermo Fisher Scientific). Size distribution and degradation were assessed using the Fragment analyzer (AATI) and High Sensitivity Large Fragment 50 kb Analysis Kit (Agilent). Purification steps were performed using AMPure PB beads (PacBio). The 32 individual samples (2 µg) were purified, then sheared at 10 kb using the Megaruptor1 system (Diagenode). Using SMRTBell template Prep Kit 1.0 and SMRTbell Barcoded Adaptater kit 8A or 8B kits (PacBio), samples (1 µg) were independently barcoded then pooled by 5 to 8. The 8 libraries were purified tree times. SMRTbell libraries were sequenced on SMRTcells on Sequel1 instrument at 6pM with 120-min preextension and 10-h or 20h movies using Sequencing Primer V4, polymerase V3, diffusion loading.

### GTWWAC methylation analysis

All methylation analyses were performed with public GMI1000 genome and annotation. Motif and methylation detection were performed using the pipeline “pbsmrtpipe.pipelines.ds_modification_motif_analysis” from PacBio SMRTLink 6.0. The default settings were used except: compute methyl fraction set as true, minimum required alignment concordance >= 80 and minimum required alignment length >= 1000.

Followed by the bioinformatics analyses of the data obtained from SMRT sequencing, methylome profiles of the 31 evolved clones were compared to the ancestral clone individually. The analysis showed the methylation profile for G**T**WW<u>A</u>C motif with a score, coverage, IPD ratio, and fraction for each sample. A score above 30 is considered significant and coverage represents the sequencing depth (higher the better). IPD ratio or interpulse duration ratio is the time required for the consequent nucleotide to bind, where the presence of methylated base increases the time required for the nucleotide addition (higher IPD ratio means a higher probability of methylation). The fraction represents the percentage of methylated bases in the genome pool at that particular position. In this experiment, the methylation or hemimethylation of a particular position is considered significant when the fraction is greater than or equal to 0.50 (represents at least 50% of the sequences are methylated at that particular position in the whole genome pool) in addition to the score above 30.

### MSRE-qPCR

The MSRE-qPCR (Methylation Sensitive Restriction Enzyme – quantitative PCR) approach was used to check the methylation profile at a specific genomic region (Krygier et al. 2016). The protocol used for MSRE-qPCR derived from Payelleville et al. (2019). Genomic DNA was extracted from bacterial cells grown in the same culture condition (synthetic MP medium with glutamine) and at the same growth stage (beginning of stationary phase) used for SMRT-seq. Genomic DNA extraction and purification was performed using the Genomic DNA Purification Kit from Promega. First, in order to generate numerous linear DNA fragments, 400 ng of genomic DNA was digested by *Eco*RI (0.25U in a total volume of 20µL) for one night at 37°C followed by an enzyme inactivation step (20 min at 65°C). Then, 8 µl of *Eco*RI-digested-DNA was digested by *Hpy166*II (0.25U in a total volume of 20 µL) for one night at 37°C followed by an enzyme inactivation step (20 min at 65°C). The *Hpy166*II restriction enzyme digests only unmethylated GTNNAC sites. A qPCR amplification was then performed on 2 µl of 10^-5^ diluted DNA in a total volume of 7 µl containing 3.6 µl of Master mix Takyon SYBR Green I and 0.5 mM of each primer. Primers used for MSRE-qPCR are described in supplementary Table S1. The qPCR amplification was performed using the LightCycler 480 II (Roche) and the following program ; 3 min of denaturation at 95°C, and 45 cycles of denaturation 10 sec at 95°C and primer annealing 45 sec at 65°C. Detection of an amplicon revealed that no digestion occurred and that the region was methylated, while non amplification revealed that the region was unmethylated and digested. The mAG4 mutant (GMI1000 deleted from the RSc1982 MTase, targeting GTWWAC motifs; see mutant construction below) was used as a non-methylated control at GTWWAC motifs (a negative control for qPCR amplification). *Eco*RI digested DNA diluted 10^-5^ times was used as a positive control for qPCR amplification.

Raw data from qPCR experiments were analyzed using the 2^-ΔΔCt^ method to perform a relative quantification (Livak and Schmittgen 2001). This method was used to relate the PCR signal of the MSRE digested DNA to the PCR signal of the *EcoR*I digested DNA. Ct values obtained with MSRE-digested DNA were first normalized with Ct values obtained with *EcoR*I-digested DNA (ΔCt = Ct_MSRE-DNA_ – Ct_EcoRI-DNA_). This ΔCt value was then normalized with the ΔCt value obtained with GMI1000 DNA (ΔΔCt = ΔCt_evolved.clone_ – ΔCt_GMI1000_). This ΔΔCt value was then normalized by the amplification efficiency coefficient of the target and reference DNAs. Here, we estimated that the two DNAs had the same amplification efficiency and close to one. Therefore, the amount of target, normalized to the reference and relative to the calibrator, was given by the 2^-ΔΔCt^ value (Livak and Schmittgen 2001). Three biological replicates (DNA extracted from independent bacterial cultures) and three technical replicates (three qPCR experiments per DNA sample) were performed. The 2^-ΔΔCt^ values were compared using the Wilcoxon non-parametric test with the R software.

### Construction of mutants

The mAG4 mutant (GMI1000 deleted from the RSc1982 MTase gene) was constructed using a SacB protocol as described in Gopalan-Nair et al. (2020). At the end of the protocol, gene deletion was checked by PCR on colonies that were resistant to sucrose and sensitive to kanamycine (plasmid lost with second recombination event).

Point mutations of the two GTWWAC motifs, changing the T by a C, upstream the RSp0338 gene were performed using the gene replacement method with the pK18 plasmid containing the *sacB* counter-selectable marker, as previously described (Gopalan-Nair et al. 2020). All the point mutations were performed on both the ancestral clone and the Mar26b2 evolved clone. All mutants were tagged with the fluorescent reporters mCherry or GFP as previously described (Perrier et al. 2019). The primers used for the construction of mutants are listed in Table S1.

### RT-qPCR analysis

The RT-qPCR (Reverse Transcription – quantitative PCR) approach was used to quantify the expression of the *epsR* gene in the ancestral GMI1000 clone, the evolved clones and the GTWWAC-epsR mutants. The protocol used for RT-qPCR derived from Perrier et al. (2016). Total RNA were isolated using TRIzol Reagent (life technologies) followed by RNeasy MiniElute Cleanup Kit (Qiagen). To avoid contamination by genomic DNA each sample was treated with the TURBO DNA-free Kit (life technologies). The reverse transcription was performed on 1 µg of total RNA using the Transcriptor Reverse Transcriptase (Roche) with random hexanucleotides primers. Quantitative PCRs were performed on a Roche LightCycler480 using The LightCycler® 480 SYBR Green I Master (Roche). Cycling conditions were as follows: 95°C for 5 min, 45 cycles at 95°C for 15 s, 60°C for 20 s and 72°C for 20 s. The specificity of each amplicon was validated with a fusion cycle. The efficiency of amplification was tested with dilution game and calculated using -1+10^1/slope^ formula. The expression of *epsR* was normalized using the geometric average of three selected reference genes (RSc0403, RSc0368 and RSp0272) for each sample and calculated using the 2^-ΔΔCt^ method (Livak and Schmittgen 2001; Rao et al. 2013). All kit and reagents were used following the manufacturer’s recommendations. The primer sets used in the experiments are listed in Table S1.

### Bacterial competition assay and serial passage experiment

The bacterial competitive assay was performed as previously described (Perrier et al. 2019). Briefly, 10 µl of the mixed inoculum, containing the GFP and mCherry clones in equal proportion at a 10^6^ CFU/ml concentration, was injected into the stem of tomato cv. Marmande, 1 cm above the cotyledons. Bacteria were recovered from the plant stem as soon as the first wilting symptoms appeared (3-5 days after inoculation) as previously described (Guidot et al. 2014).

Four serial passage experiments (SPE) into the stem of tomato cv. Marmande were performed. At each SPE, serial dilutions of the recovered bacterial suspension were conducted. Ten microliter of the 10^-3^ dilution was directly injected into the stem of a healthy plant and 50 µl of the 10^-4^ and 10^-6^ dilutions were plated on BG complete medium without triphenyltetrazolium chloride using an automatic spiral plater (easySpiral, Interscience, France). Green and red colonies were visualized and enumerated using a fluorescence stereo zoom microscope (Axio Zoom.V16, ZEISS, Germany). A competitive index (CI) was calculated at each SPE as the ratio of the two clones obtained from the plant stem (output) divided by the ratio in the inoculum (input) (Macho et al. 2010). A total of seven replicates were performed for each competition assay. Differences between mean CI values were tested using a Wilcoxon test performed in the R statistical software.

## Results

### Defining the 6mA methylation profile of strain GMI1000

In order to detect potential changes in the methylation profile of evolved clones, we first established the 6mA modification sites in the wild-type ancestor GMI1000. Methylation of 6mA type at the GTWWAC motif was investigated using SMRT-seq technology. In order to limit the number of cells in division and avoid a bias towards hemimethylated marks, genomic DNA was prepared from bacterial cells collected at the beginning of stationary phase. Growth was performed in synthetic medium with glutamine to mimic xylemic environment of the plant, glutamine being the main compound of xylem sap in most plant species (Baroukh et al. 2022).

A total of 392 GTWWAC motifs are present on the GMI1000 genome. In our culture and growth phase conditions and according to SMRT-seq data, 10 GTWWAC motifs were detected unmethylated in the GMI1000 genome, four on the chromosome and six on the megaplasmid (Table 2 and Supplemental Table S2). Eight of these unmethylated motifs were located in the upstream region of a gene, thus potentially affecting gene expression. This specifically concerned the RSc0958 gene encoding a type VI secretion system tip VgrG family protein, the *epsR* gene (two motifs) encoding the negative regulator of exopolysaccharide production (Chapman and Kao 1998) and the *efe* gene encoding the Ethylene-forming enzyme (Valls et al. 2006). We also identified 9 motifs that were hemimethylated (DNA methylation of either strand – or strand +) in the GMI1000 genome (Table 2 and Supplemental Table S2).

**Table 2.**
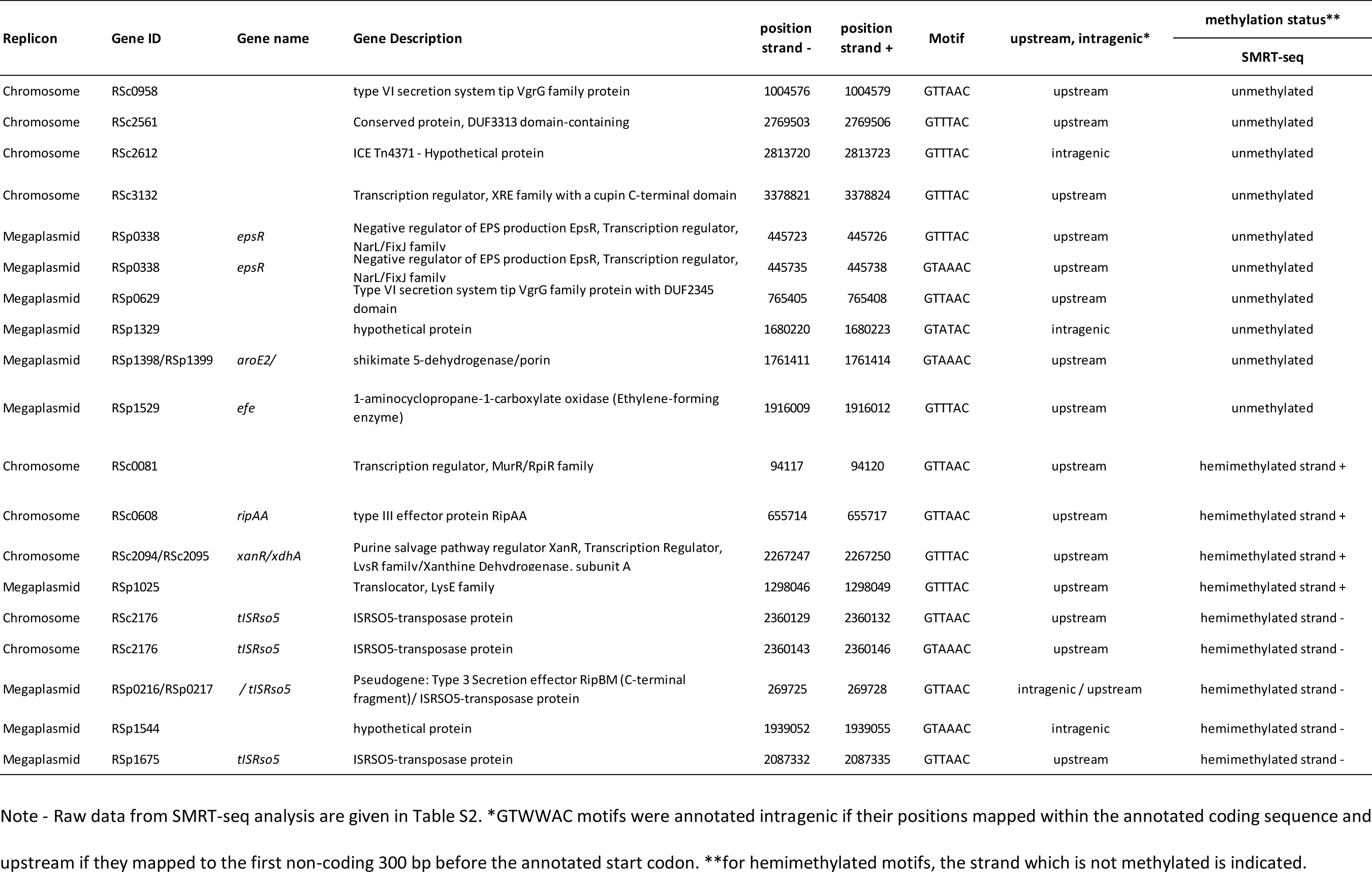
Genomic regions of the GMI1000 strain of *Ralstonia solanacearum* with a GTWWAC motif detected unmethylated or hemimethylated at the beginning of the stationary phase during growth in synthetic medium with glutamine 10mM according to SMRT-seq data

| Replicon | Gene ID | Gene name | Gene Description | position<br>strand - | position<br>strand + | Motif | upstream, intragenic* | methylation status** |
| --- | --- | --- | --- | --- | --- | --- | --- | --- |
|  |  |  |  |  |  |  |  | SMRT-seq |
| Chromosome | RSc0958 |  | type VI secretion system tip VgrG family protein | 1004576 | 1004579 | GTTAAC | upstream | unmethylated |
| Chromosome | RSc2561 |  | Conserved protein, DUF3313 domain-containing | 2769503 | 2769506 | GTTTAC | upstream | unmethylated |
| Chromosome | RSc2612 |  | ICE Tn4371 - Hypothetical protein | 2813720 | 2813723 | GTTTAC | intragenic | unmethylated |
| Chromosome | RSc3132 |  | Transcription regulator, XRE family with a cupin C-terminal domain | 3378821 | 3378824 | GTTTAC | upstream | unmethylated |
| Megaplasmid | RSp0338 | *epsR* | Negative regulator of EPS production EpsR, Transcription regulator, NarL/FixJ family | 445723 | 445726 | GTTTAC | upstream | unmethylated |
| Megaplasmid | RSp0338 | *epsR* | Negative regulator of EPS production EpsR, Transcription regulator, NarL/FixJ family | 445735 | 445738 | GTAAAC | upstream | unmethylated |
| Megaplasmid | RSp0629 |  | Type VI secretion system tip VgrG family protein with DUF2345 domain | 765405 | 765408 | GTTAAC | upstream | unmethylated |
| Megaplasmid | RSp1329 |  | hypothetical protein | 1680220 | 1680223 | GTATAC | intragenic | unmethylated |
| Megaplasmid | RSp1398/RSp1399 | *aroE2/* | shikimate 5-dehydrogenase/porin | 1761411 | 1761414 | GTAAAC | upstream | unmethylated |
| Megaplasmid | RSp1529 | *efe* | 1-aminocyclopropane-1-carboxylate oxidase (Ethylene-forming enzyme) | 1916009 | 1916012 | GTTTAC | upstream | unmethylated |
| Chromosome | RSc0081 |  | Transcription regulator, MurR/RpiR family | 94117 | 94120 | GTTAAC | upstream | hemimethylated strand + |
| Chromosome | RSc0608 | *ripAA* | type III effector protein RipAA | 655714 | 655717 | GTTAAC | upstream | hemimethylated strand + |
| Chromosome | RSc2094/RSc2095 | *xanR/xdhA* | Purine salvage pathway regulator XanR, Transcription Regulator, LysR family/Xanthine Dehydrogenase, subunit A | 2267247 | 2267250 | GTTTAC | upstream | hemimethylated strand + |
| Megaplasmid | RSp1025 |  | Translocator, LysE family | 1298046 | 1298049 | GTTTAC | upstream | hemimethylated strand + |
| Chromosome | RSc2176 | *tISRso5* | ISRSO5-transposase protein | 2360129 | 2360132 | GTTAAC | upstream | hemimethylated strand - |
| Chromosome | RSc2176 | *tISRso5* | ISRSO5-transposase protein | 2360143 | 2360146 | GTAAAC | upstream | hemimethylated strand - |
| Megaplasmid | RSp0216/RSp0217 | */ tISRso5* | Pseudogene: Type 3 Secretion effector RipBM (C-terminal fragment)/ ISRSO5-transposase protein | 269725 | 269728 | GTTAAC | intragenic / upstream | hemimethylated strand - |
| Megaplasmid | RSp1544 |  | hypothetical protein | 1939052 | 1939055 | GTAAAC | intragenic | hemimethylated strand - |
| Megaplasmid | RSp1675 | *tISRso5* | ISRSO5-transposase protein | 2087332 | 2087335 | GTTAAC | upstream | hemimethylated strand - |

Note - Raw data from SMRT-seq analysis are given in Table S2. \*GTWWAC motifs were annotated intragenic if their positions mapped within the annotated coding sequence and upstream if they mapped to the first non-coding 300 bp before the annotated start codon. \*\*for hemimethylated motifs, the strand which is not methylated is indicated.

### Mapping differential methylated sites between the ancestral and evolved clones with SMRT-sequencing

A total of 31 evolved clones derived from strain GMI1000 after experimental evolution in 5 different host plants over 300 generations were investigated. All clones but one exhibited a better fitness than their ancestral clone in their experimental host according to competition experiments (CI>1). Only the clone Zeb26d1 recovered from eggplant Zebrina had a CI not significantly different from one and was used as a control. An average of 1.2 (min 0 ; max 3) genomic polymorphisms were detected in these 31 evolved clones (Gopalan-Nair et al. 2023) (Table 1).

SMRT-seq data from the 31 evolved clones were investigated for methylome analysis, in similar conditions as for the ancestral clone. Comparison of the methylation marks on the adenine of the GTWW<u>A</u>C motifs between the ancestral clone and the 31 evolved clones revealed a list of 50 DMSs. This list included 30 DMSs at one DNA strand (hemimethylated region) and 10 DMSs at both DNA strands (Tables 3a and 3b).

**Table 3a.**
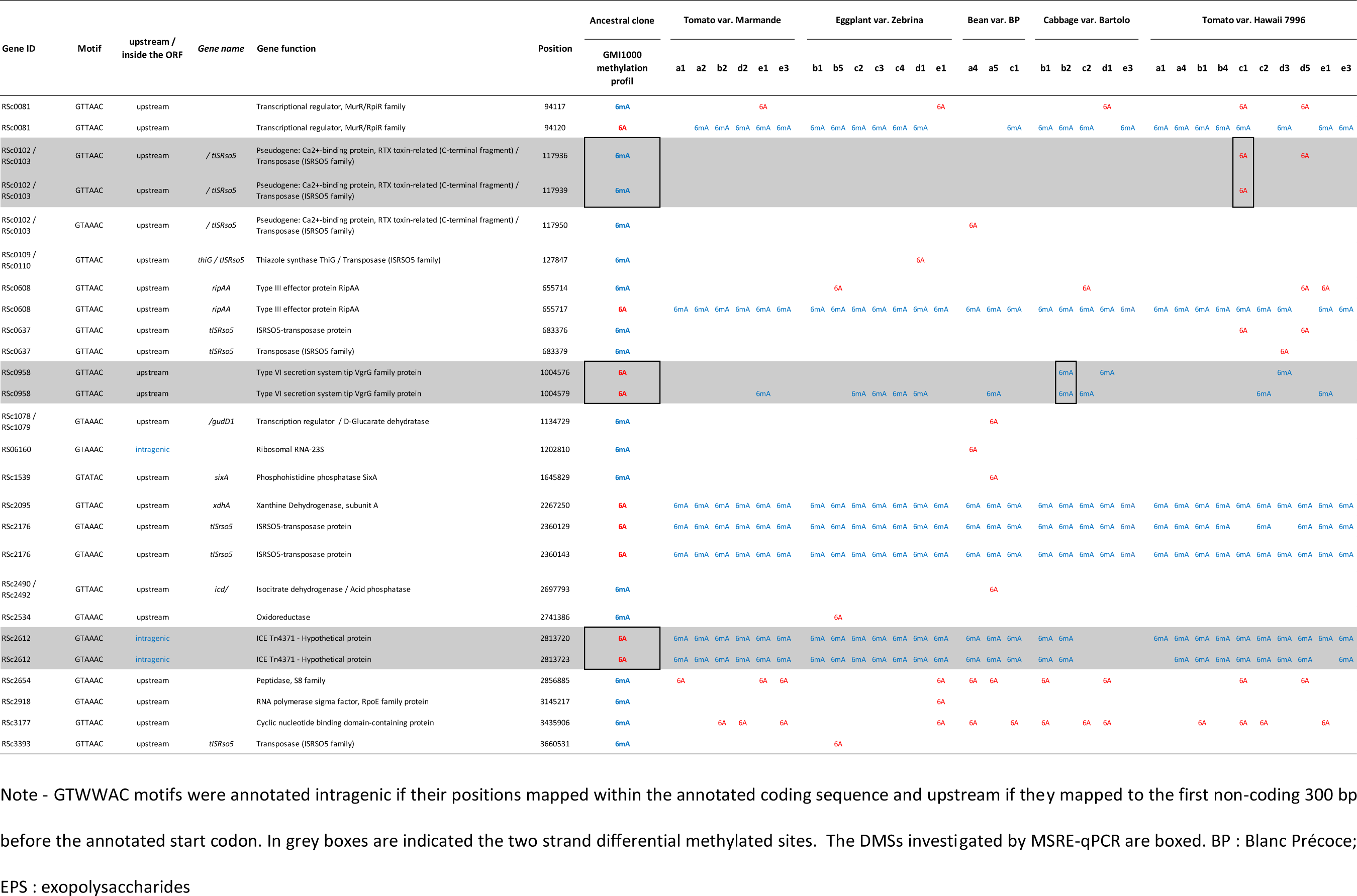
Differential methylated sites on the chromosome between the ancestral clone and the clones evolved on five different plant species.

**Table 3b.**
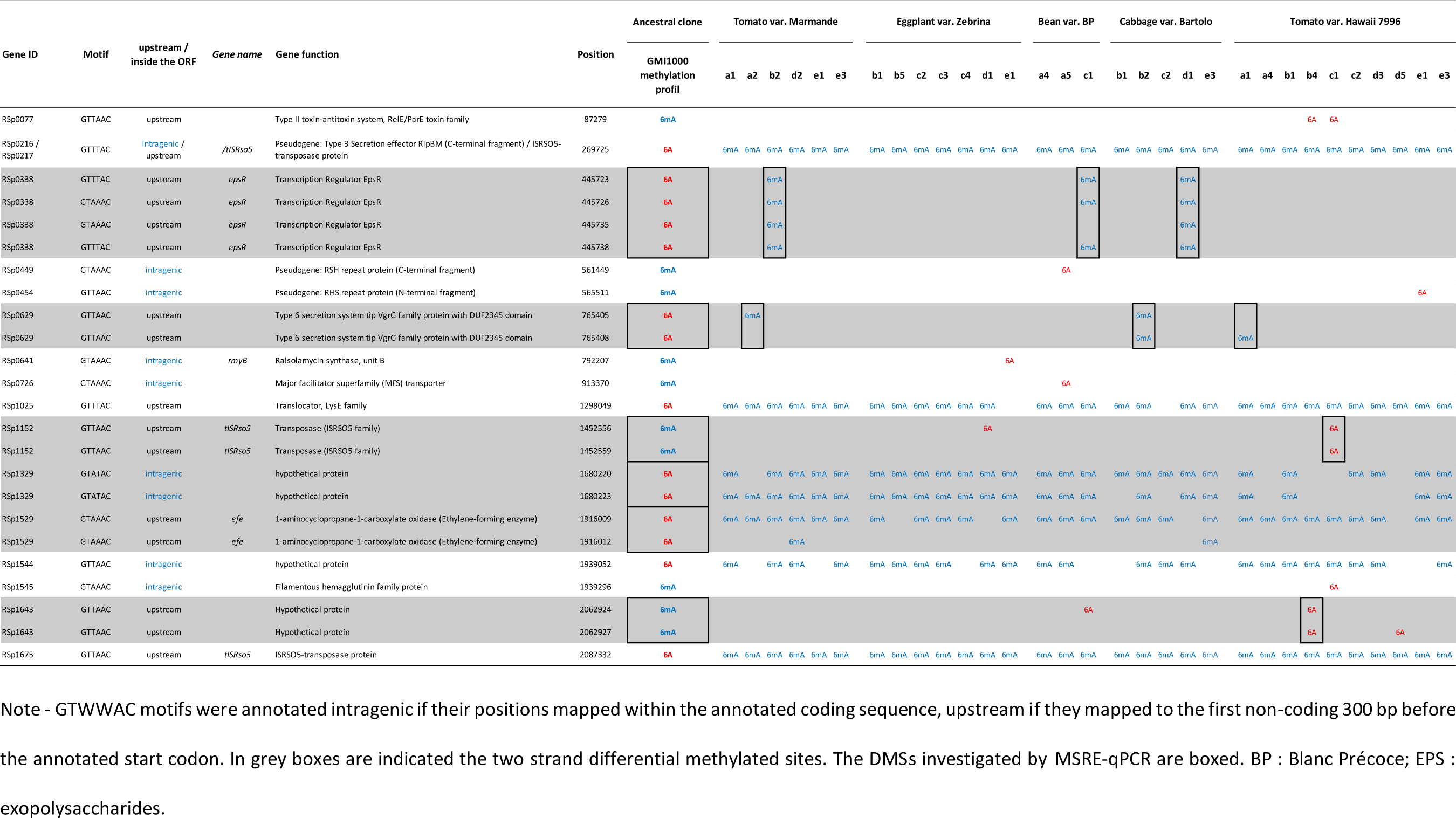
Differential methylated sites on the megaplasmid between the ancestral clone and the clones evolved on five different plant species.

### Characteristics of the DMSs

Methylome comparison between the ancestral clone and the 31 evolved clones revealed between 12 and 21 (15.5 ± 2.2 ; mean ± standard deviation) DMSs per evolved clone (Tables 3a,3b and Supplemental Figure 1). The experimental host did not have a strong impact on the number of DMSs, with the exception that the number of DMSs detected in bean clones was significantly superior to the number of DMSs detected in eggplant Zebrina and in tomato Hawaii clones (Supplemental Figure 1).

Genomic repartition analysis of the DMSs revealed that 26 were on the chromosome (3.7 Mb) and 24 on the megaplasmid (2.1 Mb) which seems to indicate a higher frequency on the second replicon (Table 4 and Figure 1). However, the examination of the map does not reveal any specific region enriched in hypo or hypermethylation (Figure 1).

**Table 4.** General features of the differential methylated sites (DMSs) identified in evolved clones from five different plant species.

|  | Genome<br>(5.8 Mbp) | Chromosome<br>(3.7 Mbp) | Megaplasmid<br>(2.1 Mbp) |
| --- | --- | --- | --- |
| **nbr of DMSs** | **50** | **26** | **24** |
| *DMS frequency / Mbp* | *8,62* | *7,03* | *11,43* |
| **nbr of genes affected by DMSs** | **39** | **22** | **17** |
| **nbr of DMSs in gene promoter region** | **39** | **23** | **16** |
| *% DMSs in gene promoter regions* | *78* | *88* | *67* |
| **nbr of DMSs affecting transposable elements** | **13** | **9** | **4** |
| *% transposable elements/nbr of DMSs in gene promoter regions* | *33* | *39* | *25* |
| **nbr of DMSs affecting virulence determinants** | **12** | **4** | **8** |
| *% virulence determinants/nbr of DMSs in gene promoter regions* | *31* | *17* | *50* |

Note - \*Gene promoter region was defined as the first non-coding 300 bp region before the annotated start codon of the gene. Mbp : million of base pairs

**Figure 1.**
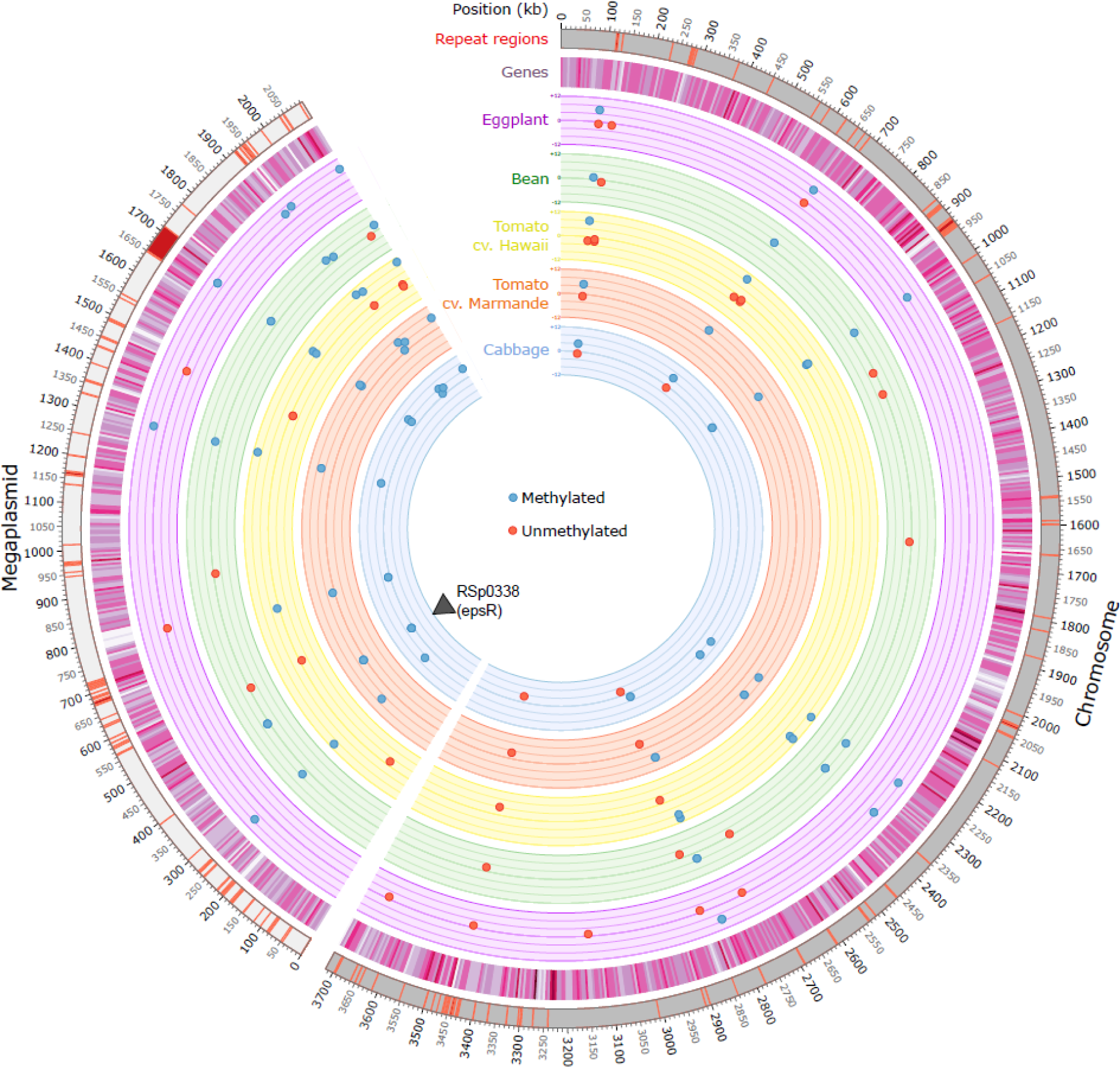
Circos plot highlighting the genomic repartition of the sites differentially methylated (DMS) between an ancestral clone and 31 clones evolved during 300 generations on five different plant species. Newly methylated sites are indicated in blue and unmethylated sites in red. A total of 31 evolved clones were investigated; 6 evolved on tomato cv. Marmande, 7 on eggplant, 3 on bean and 10 on tomato cv. Hawaii. The number of clones targeted by a DMS is indicated on the scale varying between 0 and 12 for each plant species. The black triangle indicates the position of the RSp0338 gene.

DMSs can be classified as intragenic (position within a coding sequence), or intergenic either at the 5’ (upstream) or 3’ (downstream) position of a gene. Due to the existence of divergent promoters, a DMS at the 5’ position can potentially affect two genes, which explains why the number of genes potentially affected by these DMSs (39 genes) is slightly different from the number of DMSs (Table 4). Clearly, the number of DMSs positioned in a gene promoter region (defined as less than 300 nucleotides from the start codon) of the 39 affected genes is predominant (78%). Interestingly, one regulatory gene (RSp0338) has two GTWWAC motifs in its promoter region, both differentially methylated on both DNA strands (Table 3b). An examination of the list of the DMSs affecting promoter regions revealed an overabundance of genes encoding transposable elements (33%) and genes closely or remotely associated with virulence (*epsR*, *efe*, the type III effector *ripAA* and VGR-related proteins linked to the type VI secretion system) (31%) (Table 4).

### Differential methylation does not appear to be correlated with differential gene expression

Transcriptome analyses for the ancestral clone and the 31 evolved clones were performed by RNA sequencing in our previous work (Gopalan-Nair et al. 2023). Table 5a gives a summary of the relative gene expression in the experimentally evolved clones compared to the ancestral clone for each of the 39 genes targeted by a DMS. A Fisher exact test was used to determine whether there was an association between differential methylation and differential gene expression. This analysis revealed a significant correlation only for the RSp0338 gene (Table 5b).

**Table 5a.**
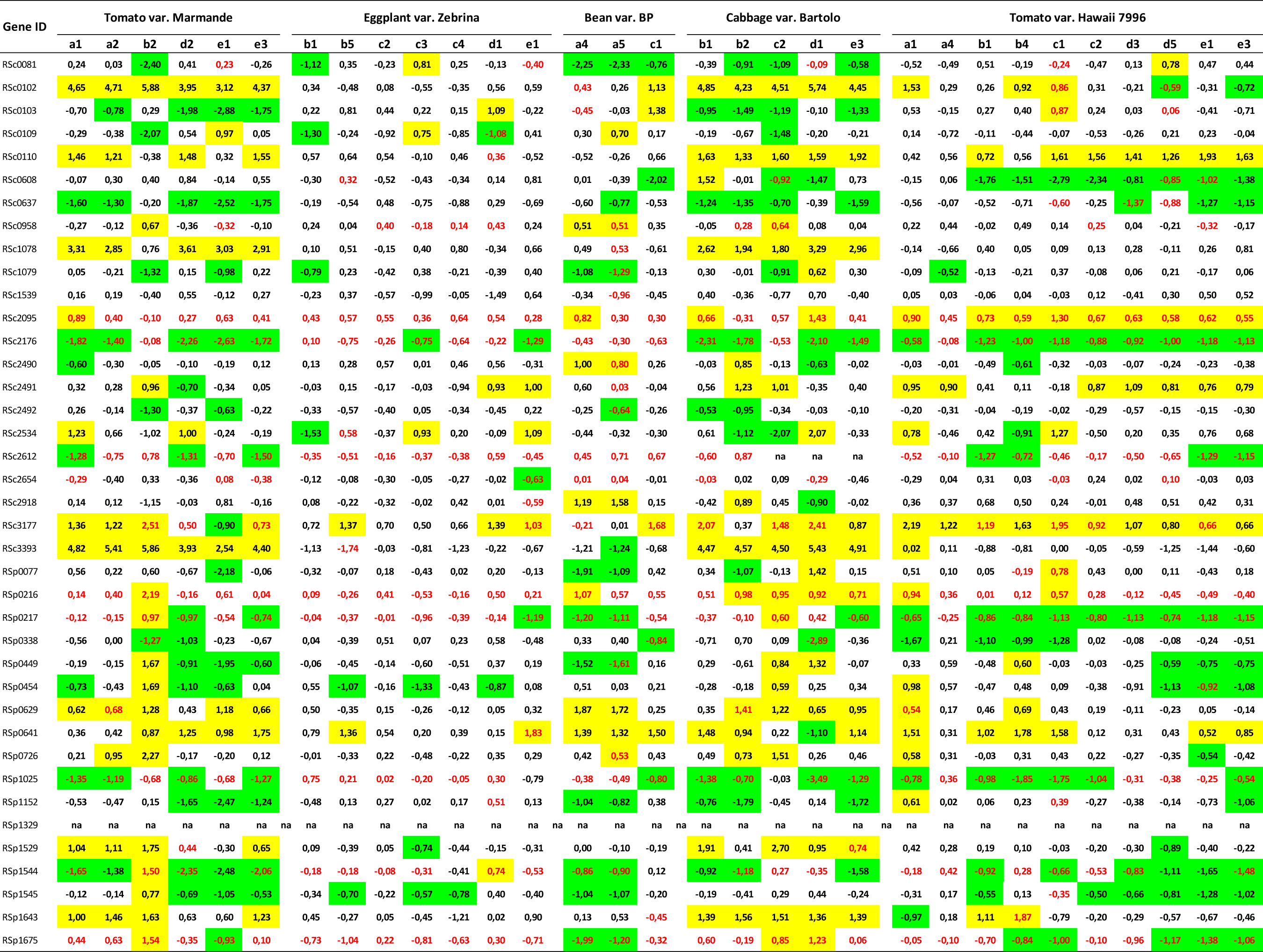
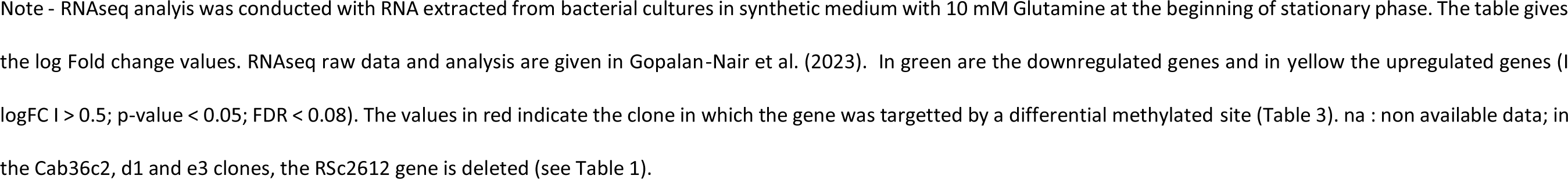
Relative gene expression in the experimentally evolved clone compared to the ancestral clone for each gene targetted by a differential methylated site.

**Table 5b.** Association analysis between differential methylation and differential gene expression between the ancestral clone and the experimentally evolved clones.

| Gene ID | DMS-DEG | DMS-non DEG | non DMS-DEG | non DMS-non DEG | p-value (Fisher exact Test) |
| --- | --- | --- | --- | --- | --- |
| RSc0081 | 0 | 4 | 10 | 17 | 0,2770 |
| RSc0102 | 2 | 1 | 15 | 13 | 1,0000 |
| RSc0103 | 1 | 2 | 10 | 18 | 1,0000 |
| RSc0109 | 1 | 0 | 6 | 24 | 0,2258 |
| RSc0110 | 0 | 1 | 16 | 14 | 0,4839 |
| RSc0608 | 3 | 1 | 9 | 18 | 0,2718 |
| RSc0637 | 1 | 2 | 12 | 16 | 1,0000 |
| RSc0958 | 2 | 8 | 2 | 19 | 0,5773 |
| RSc1078 | 0 | 1 | 10 | 20 | 1,0000 |
| RSc1079 | 1 | 0 | 7 | 23 | 0,2581 |
| RSc1539 | 0 | 1 | 0 | 30 | 1,0000 |
| RSc2095 | 13 | 18 | 0 | 0 | 1,0000 |
| RSc2176 | 20 | 11 | 0 | 0 | 1,0000 |
| RSc2490 | 1 | 0 | 5 | 25 | 0,1935 |
| RSc2491 | 0 | 1 | 13 | 17 | 1,0000 |
| RSc2492 | 1 | 0 | 3 | 27 | 0,1290 |
| RSc2534 | 0 | 1 | 11 | 19 | 1,0000 |
| RSc2612 | 7 | 21 | 0 | 0 | 1,0000 |
| RSc2654 | 1 | 9 | 0 | 21 | 0,3226 |
| RSc2918 | 0 | 1 | 4 | 26 | 1,0000 |
| RSc3177 | 11 | 2 | 12 | 6 | 0,4120 |
| RSc3393 | 0 | 1 | 12 | 18 | 1,0000 |
| RSp0077 | 1 | 1 | 5 | 24 | 0,3548 |
| RSp0216 | 8 | 23 | 0 | 0 | 1,0000 |
| RSp0217 | 17 | 14 | 0 | 0 | 1,0000 |
| RSp0338 | 3 | 0 | 5 | 23 | 0,0125 |
| RSp0449 | 1 | 0 | 11 | 19 | 0,3871 |
| RSp0454 | 1 | 0 | 11 | 19 | 0,3871 |
| RSp0629 | 3 | 0 | 10 | 18 | 0,0636 |
| RSp0641 | 1 | 0 | 18 | 12 | 1,0000 |
| RSp0726 | 1 | 0 | 6 | 24 | 0,2258 |
| RSp1025 | 15 | 14 | 0 | 2 | 0,4839 |
| RSp1152 | 0 | 2 | 10 | 19 | 1,0000 |
| RSp1329 | na | na | na | na | na |
| RSp1529 | 1 | 1 | 9 | 20 | 1,0000 |
| RSp1544 | 12 | 11 | 6 | 2 | 0,412 |
| RSp1545 | 0 | 1 | 15 | 15 | 1,0000 |
| RSp1643 | 1 | 1 | 11 | 18 | 1,0000 |
| RSp1675 | 11 | 20 | 0 | 0 | 1,0000 |

Note - For each gene targetted by a differentially methylated site (DMS) in at least one experimentally evolved clone, the table gives the number of clones in which the gene is both differentially methylated and differentially expressed (DMS-DEG (Differentially espressed gene)), differentially methylated but not differentially expressed (DMS-non DEG), not differentially methylated but differentially expressed (non DMS-DEG) and neither differentially methylated nor expressed (non DMS - non DEG). A fisher exact test was used to determine whether there was an association between methylation and gene expression.

Down regulation of the RSp0338 gene in the Mar26b2, Bean26c1 and Cab36d1 clones compared to the ancestral GMI1000 clone was investigated using a RT-qPCR approach. This analysis showed that the RSp0338 gene is down-regulated in the three investigated evolved clones compared to the ancestral GMI1000 clone (Figure 2).

**Figure 2.**
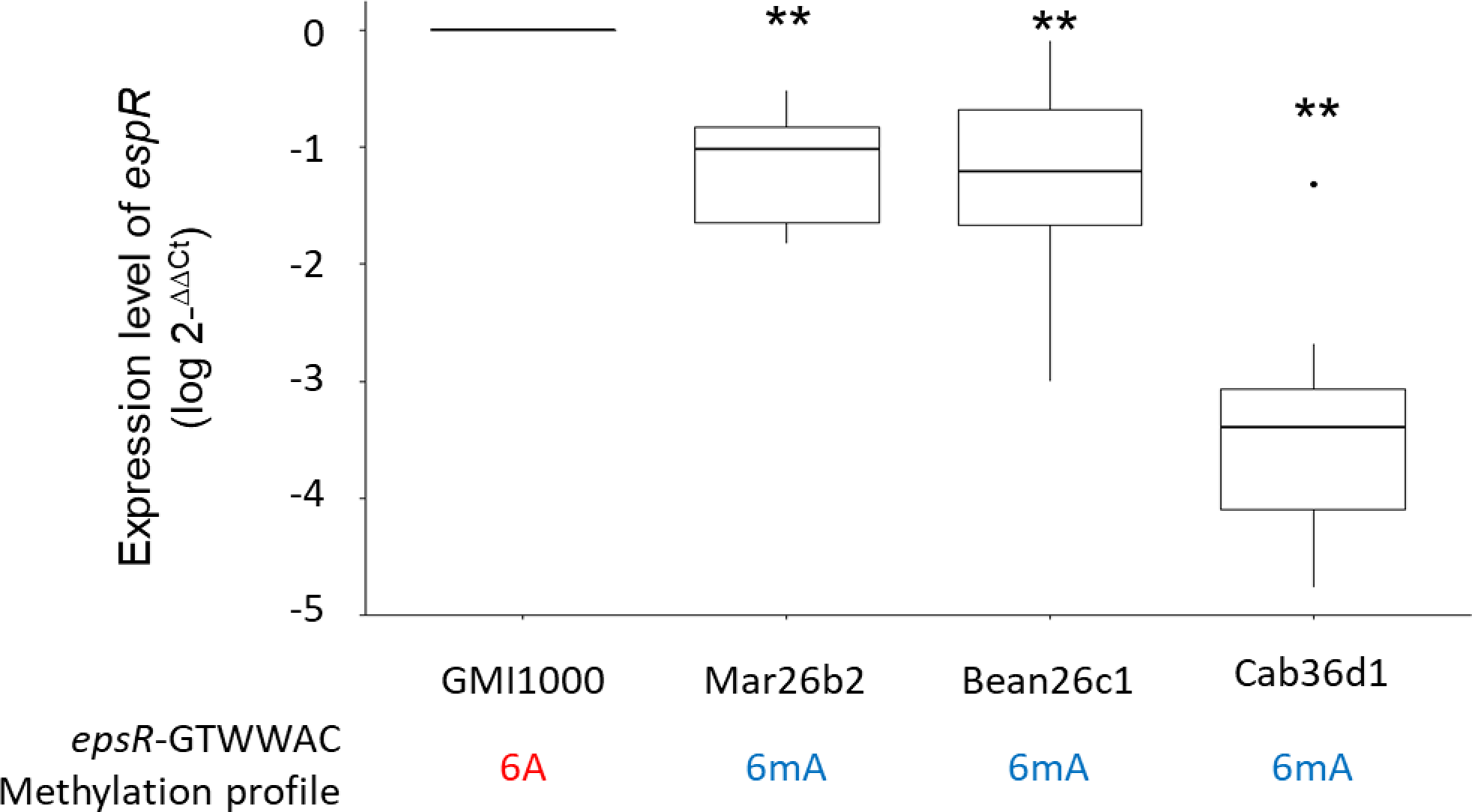
Relative expression level of RSp0338 gene between GMI1000 and evolved clones. Expression level of RSp0338 was determined during growth in synthetic medium supplemented with Glutamine 10 mM at the beginning of stationary phase, using a RT-qPCR approach. The methylation profile of the GTWWAC motifs in the upstream region of epsR is indicated for each investigated clone. Three technical and three biological replicates were performed. Data were normalized using the 2-ΔΔCt calculation method (Livak and Schmittgen 2001). (Wilcoxon test, ** p value < 0,01).

### Assessment of methylation status through the MSRE-qPCR approach

We used MSRE-qPCR (Methylation Sensitive Restriction Enzyme-quantitative PCR) assay as an alternative approach to assess the methylation status of DMSs identified by SMRT-seq. Briefly, MSRE-qPCR is based on extensive digestion of genomic DNA with methylation-sensitive restriction enzyme (MSRE) followed by quantitative PCR amplification of the target gene (Krygier et al. 2016). With this method, we could only test two-strand-DMSs, but not hemimethylated sites. Genomic DNA was prepared in similar conditions as for SMRT-seq.

The MSRE-qPCR approach was first used to assess the methylation status of the GMI1000 strain for three motifs that were detected methylated on both DNA strands for a majority of the evolved clones but not methylated in the ancestral clone according to SMRT-seq. These three motifs were associated with the RSc2612, RSc1329 and RSc1529 genes (Table 3). According to the MSRE-qPCR results, the ancestral clone was found methylated such as the evolved clones (Figure 3).

**Figure 3.**
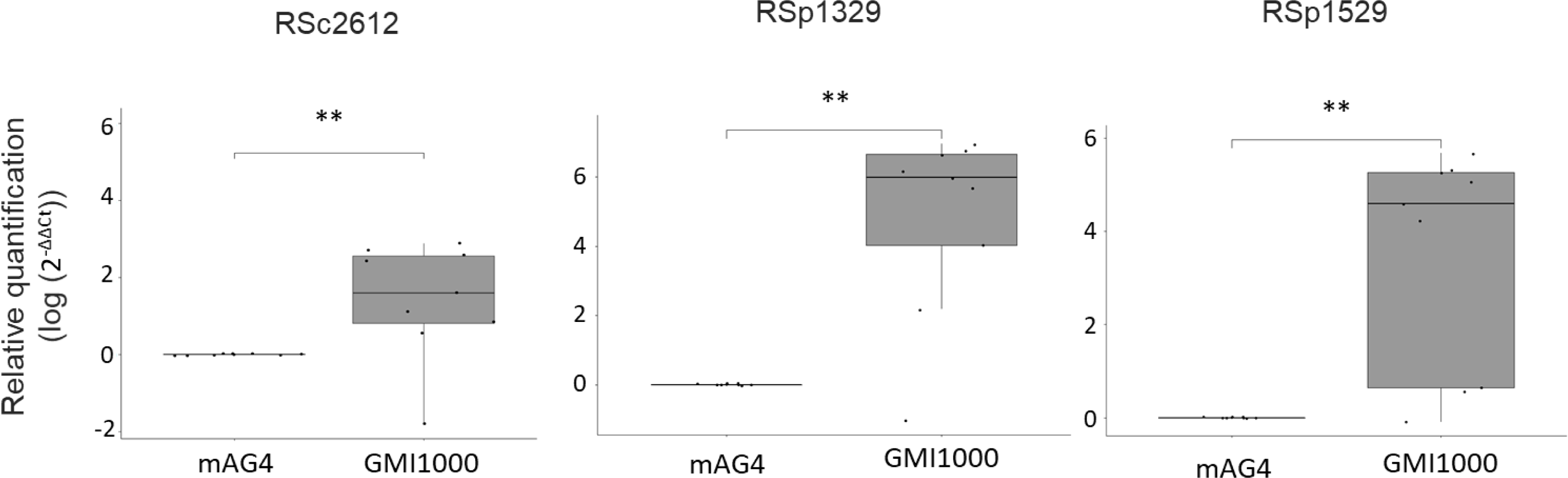
MSRE-qPCR results for analysis of methylation status of GTWWAC motifs at the RSc2612, RSp1329 and RSp1529 genes in the ancestral GMI1000 clone. The methylation profile of the GTWWAC motifs at the RSc2612, RSp1329 and RSp1529 genes for the ancestral GMI1000 clone was investigated using a MSRE-qPCR approach (Methylation Sensitive Restriction Enzyme-quantitative PCR). Bacterial cells were grown in synthetic medium with glutamine 10 mM and DNA was recovered at the beginning of stationary phase. The mAG4 mutant (GMI1000 deleted from the RSc1982 MTase, targeting GTWWAC motifs) was used as a non-methylated control at GTWWAC motifs. The graphs represent a relative quantification using the 2-ΔΔCt method compared to the mAG4 mutant. Detection of an amplicon revealed that no digestion occurred and that the region was methylated, while non amplification revealed that the region was non-methylated and digested. 2-ΔΔCt values were compared between the ancestral clone and mAG4 mutant using a Wilcoxon test; ** p-value < 0.01.

We then used MSRE-qPCR to investigate the methylation status of seven motifs found differentially methylated according to SMRT-seq. These seven motifs included one motif in the divergent promoter region of both RSc0102 and RSc0103, two motifs upstream of RSp0338 and one motif upstream of RSc0958, RSp0629, RSp1152 and RSp1643 (Tables 3a and 3b). MSRE-qPCR analysis was conducted on both GMI1000 DNA and DNA from the evolved clones in which the two-strands-DMSs were found (Table 3a and 3b). For the RSp0338 and RSp0629 genes, we also included in the analysis DNA from three evolved clones in which differential hemimethylation was detected. This concerned the Bean26c1 clone for RSp0338 and Mar26a2 and Haw35a1 clones for RSp0629. According to the MSRE-qPCR results, the GTWWAC motifs upstream of RSc0102/RSc0103, RSc0958, RSp0629, RSp1152 and RSp1643 were not found differentially methylated between the ancestral and the evolved clones, being fully methylated in both (Figure 4). However, the MSRE-qPCR analysis showed that the region upstream of RSp0338 was differentially methylated between GMI1000 and the three independent clones evolved on tomato var. Marmande, bean and cabbage. In agreement with SMRT-seq data, the GTWWAC motifs upstream of RSp0338 appeared not methylated in the ancestral clone but methylated in the Mar26b2, Bean26c1 and Cab36d1 evolved clones (Figure 4).

**Figure 4.**
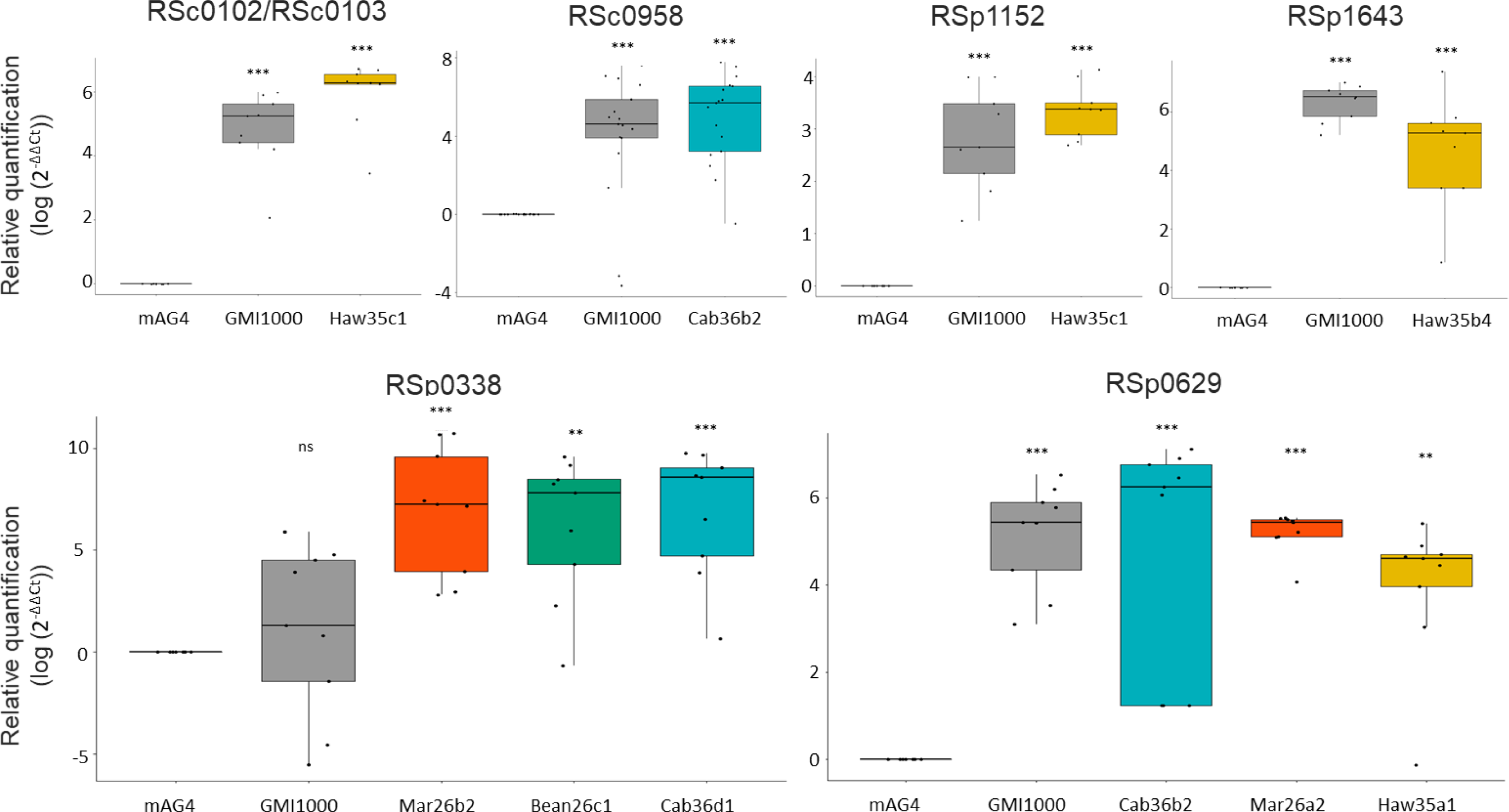
MSRE-qPCR results for analysis of methylation status of GTWWAC motifs upstream the RSc0102, RSc0958, RSp0338, RSp0629, RSp1152 and RSp1643 genes in the ancestral GMI1000 clone and the experimentally evolved clones. See Figure 3 for legend. 2-ΔΔCt values were compared between the evolved or ancestral clone and mAG4 mutant using a Wilcoxon test; ns: not significant; ** p-value < 0.01; *** p-value < 0.001.

### Methylation upstream of the RSp0338 gene appeared after only 2 passages in plant

In this part of the study, we were interested in determining at which evolution stage did the RSp0338 differential methylation arises. To answer this question, we conducted an MSRE-qPCR analysis on DNA from clones from the tomato Marmande lineage B evolved after one, two, three, four, five, ten, 14, 18 and 22 serial passages. Two clones per serial passage were investigated. The MSRE-qPCR results showed that the two GTWWAC motifs upstream RSp0338 were not methylated for the two clones recovered after one passage, as for the ancestral clone. However, the motifs were already methylated in one of the two clones recovered after two and three passages and remain methylated in all the clones recovered in the following passages (Figure 5).

**Figure 5.**
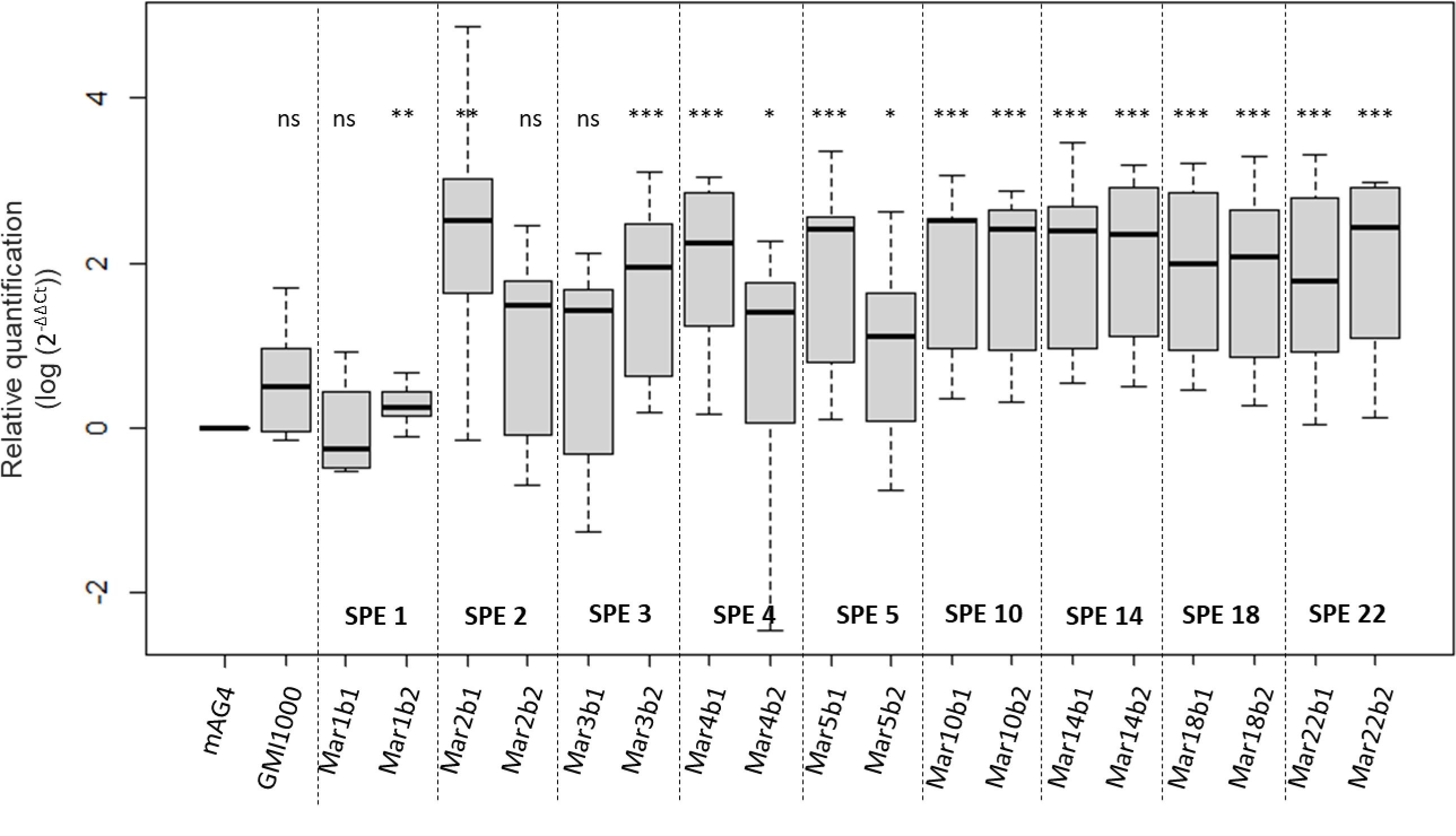
MSRE-qPCR results for chronology of methylation appearance upstream RSp0338 during experimental evolution in tomato Marmande. The methylation profile of the GTWWAC motifs upstream RSp0338 was investigated using a MSRE-qPCR approach for the ancestral GMI1000 clone and the ongoing experimentally evolved clones in tomato Marmande host. Evolved clones in tomato Marmande from lineage B were tested at different serial passaging during experimental evolution. Evolved clones are designate with MarXbx notation with X as the number of serial passage experiment (SPE) and x as the clone number. See figure 4 and 3 for legend.

### Methylation in the upstream region of RSp0338 contributes to bacterial fitness

In our previous work, we demonstrated that the Mar26b2 clone showed a fitness advantage during growth into the stem of its experimental host, tomato var. Marmande, compared to the ancestral GMI1000 clone, using a competition experiment approach (Table 1; Guidot et al. 2014).

In order to analyze the contribution of methylation in the upstream region of the RSp0338 gene in fitness gain of the Mar26b2 clone into tomato var. Marmande, we first constructed mutants of both GMI1000 and Mar26b2 strains in which the two GTWWAC motifs in the upstream region of RSp0338 were modified, so that they can no longer be methylated. The GTWWAC motifs modification was performed by introduction of a point mutation replacing the T by a C (Table 6). In a second step, we measured the impact of these mutations on the bacterial fitness into tomato var. Marmande. Our hypothesis was that the strains having a fitness advantage into tomato var. Marmande should enhance their frequency in the population after serial passage experiments (SPE) in this host. We thus conducted SPE in tomato var. Marmande starting with a mixed inoculum of the investigated clones and mutants and measured the competitive index (CI) after each passage (Figure 6A). Competition SPE with the Mar26b2 and GMI1000 clones validated the fitness advantage of the Mar26b2 clone with CI values enhancing at each passage (Figure 6B). Competition SPE with the GMI1000 mutant and GMI1000 wild-type strain showed that the CI values were not significantly different from one at each passage, thus demonstrating that point mutations of the GTWWAC motifs of the RSp0338 upstream region did not impact the fitness of the GMI1000 strain (Figure 6C). Competition SPE with the Mar26b2 clone and Mar26b2 mutant showed an increase in CI values at each passage (even if this increase was not as high as the increase observed for Mar26b2 and GMI1000 competition), thus demonstrating a fitness advantage of Mar26b2 clone compared to Mar26b2 mutant (Figure 6D). Considering that point mutations of the GTWWAC motifs of the RSp0338 upstream region did not impact the fitness (Figure 6C), these results showed a role of methylation of these GTWWAC motifs in adaptive advantage of Mar26b2 clone for growth into the stem of tomato var. Marmande.

**Table 6.**
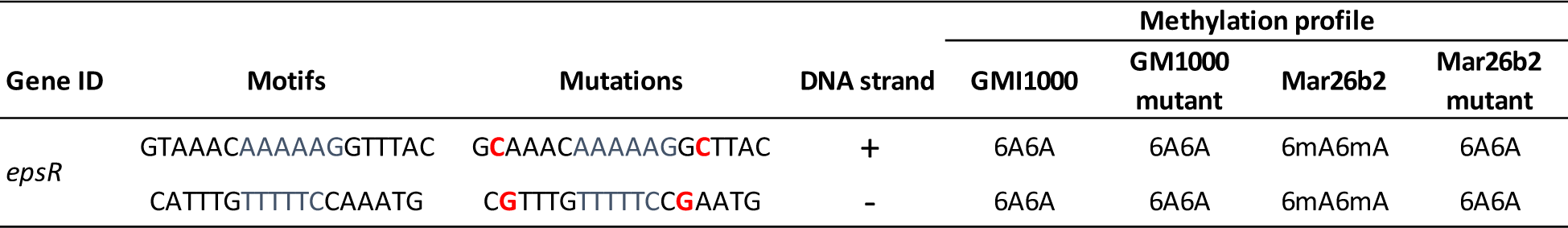
Methylation profiles of GMI1000 and Mar26b2 clones and their corresponding epsR-GTWWAC mutants at the beginning of the stationnary phase during growth in synthetic medium with glutamine 10 mM.

**Figure 6.**
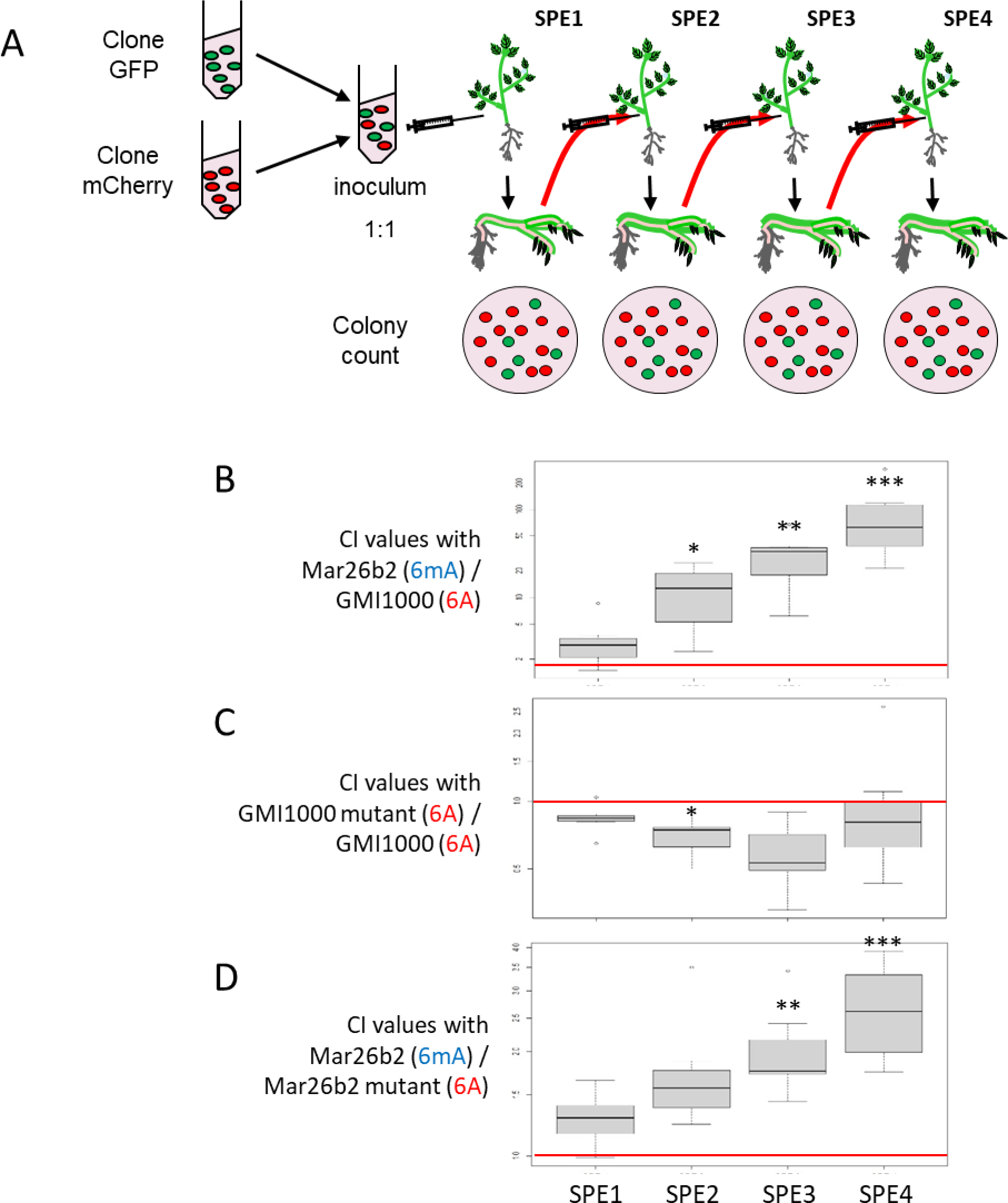
Impact of mutation of the GTWWAC motif in the upstream region of RSp0338 gene on bacterial fitness during growth into tomato var. Marmande. (A) Serial passage experiments (SPE) were conducted starting with a mixed inoculum of two clones, tagged with a GFP or mCherry marker, in the same proportion. At each passage, the competitive index (CI) between the two clones was calculated. (B) CI values of the Mar26b2 evolved clone in competition with the GMI1000 ancestral clone after 1, 2, 3 and 4 SPE. (C) CI values of the GMI1000 mutant in competition with the GMI1000 ancestral clone after 1, 2, 3 and 4 SPE. (D) CI values of the Mar26b2 evolved clone in competition with the Mar26b2 mutant after 1, 2, 3 and 4 SPE. In brackets are indicated the methylation profiles of the GTWWAC motifs in the upstream region of the RSp0338 gene for each investigated clone and mutant. The red bar highlights CI=1. Wilcoxon test, *p-value < 0.05; **p-value < 0.01; ***p-value < 0.001.

## Discussion

### DNA methylation changes during experimental adaptation of *R. solanacearum* to multihost species

In our previous works, transcriptomic analyses of experimentally adapted clones of *R. solanacearum* to various host plants revealed important variations in gene expression even in clones with no genomic alteration (Gopalan-Nair et al. 2020; Gopalan-Nair et al., 2023). Here, we investigated the methylomes of these evolved clones using SMRT-seq technology, which identified a list of 50 putative differentially methylated sites at the GTWWAC motif with a varying number of 12 to 21 DMSs per evolved clone. This list included 30 differential hemimethylated (one DNA strand) and 10 differential methylated sites (both DNA strands). In bacteria, hemimethylated DNA is produced at every round of DNA replication. This DNA modification is generally transient because the DNA methyltransferases will quickly re-methylate the majority of their target motifs. However, stable hemimethylated and unmethylated motifs have been reported in various organisms including bacteria (Payelleville et al. 2018; Sharif and Koseki 2018; Sánchez-Romero and Casadesús 2020). This phenomenon is well documented in *E. coli* and *S. typhimurium* where stable hemimethylated and unmethylated GATC sites are formed when a DNA-binding protein protects hemimethylated DNA from Dam methylase activity (Sánchez-Romero and Casadesús 2020). Differential methylation pattern on the DNA are involved in phenotypic variation by impacting gene expression through the differential affinity of some transcription factors for methylated *versus* unmethylated or hemimethylated promoters (Casadesús and Low 2013).

The MSRE-qPCR approach was used as an alternative methodology to investigate the methylation state of the 10 two-strand-DMSs detected by SMRT-seq. MSRE-qPCR appeared to be more stringent, founding only a small proportion of the differential methylated sites detected by SMRT-seq. Only one site, upstream of RSp0338, was detected between the ancestral clone and three evolved clones to be differentially methylated by using the MSRE-qPCR approach. A technical reason may explain this discrepancy, because restriction endonuclease sensitive to methylation can display various rates of cleavage depending on several parameters (time of digestion, amount of enzyme, flanking sequence…), and therefore do not always cut 100% of the DNA motifs they recognize (Roberts et al. 2015). Another possible reason for this discrepancy could be dependent on phenotypic heterogeneity, which is common in bacterial populations (Casadesús and Low 2013). This phenomenon has already been observed in populations of *R. solanacearum* GMI1000 (Perrier et al. 2019) and several mechanisms involved in the generation of phenotypic heterogeneity include epigenetic regulations (Casadesús and Low 2013; Parab et al. 2022). This could explain why different methylation states were found using either the SMRT-seq or MSRE-qPCR technologies. It should be remembered that MSRE-qPCR can only detect two-strand methylated sites, unlike SMRT-seq, but it is likely that both methods generate false positives. Nevertheless, SMRT-seq already provides a first comprehensive view of 6mA methylation profile of both ancestral and evolved clones, and the combination of the two methods has enabled us to robustly validate two differential methylation sites upstream RSp0338 between the ancestral and three evolved clones.

### DNA methylation changes rarely correlate with changes in gene expression

Among the 31 investigated evolved clones, 39 genes had a potential DMS mark. The analysis of the association between differential methylation and differential gene expression, however, revealed a significant correlation only for the RSp0338 gene. This data supports a recent analysis of the *Salmonella typhimurium* methylome and transcriptome showing that DNA methylation changes generally do not correlate with obvious changes in gene expression (Bourgeois et al. 2022).

Concerning the RSp0338 gene, the two GTWWAC motifs that were detected as differentially methylated, are located 321 bp and 309 bp upstream the start codon, thus potentially affecting the promoter region. The correlation between differential methylation and differential expression of the RSp0338 gene suggested an epigenetic regulation, a phenomenon reported in prokaryotes although still scarcely investigated (Payelleville and Brillard 2021). Epigenetic regulation in bacteria was reported to result from the impact of DNA methylation on the interaction of DNA-binding proteins with their cognate sites or on changes in DNA topology (Casadesús and Low 2013; Casadesús 2016; Sánchez-Romero and Casadesús 2020). Here, we provide evidence that RSp0338 is a novel example of epigenetically regulated gene in bacteria.

### Why adapt through methylation?

Epigenetic mutations are known to occur at a faster rate than genetic mutation (van der Graaf et al. 2015; Hu et al. 2019). The novel methylation state of the RSp0338 promoter appeared very quickly in the experimental evolution since they are detected from the first two or three serial passages on the hosts. We hypothesize that such fast epigenetic changes can allow rapid adaptation to new environmental conditions. There’s also the plausibility that epimutation is easier to generate (and especially to revert) than a genetic mutation, and that this property is therefore favorable to rapid adaptation in fluctuating environments. A question that remains unanswered is the stability of the novel methylation profile and how it will influence long-term adaptation to new environments. More and more studies report the existence of stable ‘epialleles’ that are transmitted intergenerationally and affect the phenotype of offsprings. In the same way as conventional DNA sequence-based alleles, these epialleles could be subjected to natural selection thus contributing to long-term evolutionary processes (Ashe et al. 2021). Other studies support the hypothesis of the genetic assimilation theory by which epigenetic changes could facilitate genetic mutation assimilation (Ehrenreich and Pfennig 2016; Kronholm and Collins 2016; Kronholm et al. 2017; Danchin et al. 2019; Stajic et al. 2019; Walworth et al. 2021).

### Evidence that methylation changes in RSp0338 (*epsR*) provides adaptation

Using a site-directed mutagenesis approach targeting the GTWWAC motifs that were detected differentially methylated between the evolved clones and the ancestral clone, we prevented the methylation by the MTase. An *in planta* competition experiment between the mutant and the evolved clone demonstrated that methylation of the motifs in the upstream region of the RSp0338 gene gives an adaptive advantage. To our knowledge, this is the first study showing a link between epigenetic variation and evolutionary adaptation to new environment. The involvement of epigenetic variation in environmental adaptation has been reported in several eukaryotic species (Weiner and Katz 2021; Vogt 2023). In bacteria, this has been reported for adaptation to antibiotic treatments (Ghosh et al. 2020; Muhammad et al. 2022).

The Rsp0338 gene has been characterized in the past as *epsR* (Chapman and Kao 1998), but its function remains unclear. EpsR, a putative DNA-binding protein, was shown to regulate exopolysaccharids (EPS) production in *R. solanacearum* since its overproduction strongly represses EPS synthesis but inactivation of the gene did not obviously affect EPS production (Chapman and Kao 1998; Garg et al. 2000). Based on this knowledge, it is difficult to infer a role for the decrease in *epsR* expression (as suggested by the transcriptomic data from evolved clones) linked to methylation of its promoter. Nevertheless, it is certain that *epsR* is, directly or indirectly, linked to the PhcA-dependent virulence regulation network in *R. solanacearum* (Garg et al. 2000; Genin and Denny 2012), and probably contributes to the control of EPS production or associated molecules. It should be noted that during the evolution of GMI1000 by serial passages on several host plants, alterations in another regulatory gene, *efpR*, conferring strong adaptive gains were selected and lead to multiple phenotypic changes, including significant modifications for EPS production (Perrier et al. 2016; Perrier et al. 2019). We can therefore hypothesize that the production of these surface/excreted molecules plays an important role in the phases of adaptation to the environmental conditions encountered during plant infection, and future work will need to establish their role at this level.

## Supplementary materials

**Supplemental Figure 1** Effect of the experimental host on the number of differential methylated sites (DMSs) detected in the evolved clones according to SMRT-sequencing. A. Number of DMSs in each investigated evolved clone. B. Mean number of DMSs in evolved clones for each experimental host. Different letters above the boxplot indicate a significant difference (Wilcoxon test, p.value < 0.05). Mar: Tomato var. Marmande; Zeb : Eggplant var. Zebrina; Bean : Bean var. Blanc précoce; Cab : Cabbage var. Bartolo; Haw: Tomato var. Hawaii 7996.

**Table S1** List of primers used in this study

**Table S2** Genomic regions of the GMI1000 strain of Ralstonia solanacearum with a GTWWAC motif and methylation status at the beginning of the stationary phase during growth in minimal medium with glutamine 10mM

