## Supplementary material for "Evidence for increased fitness of a plant pathogen conferred by epigenetic variation": Figure S1

Gopalan-Nair *et al.*, Sup-figures

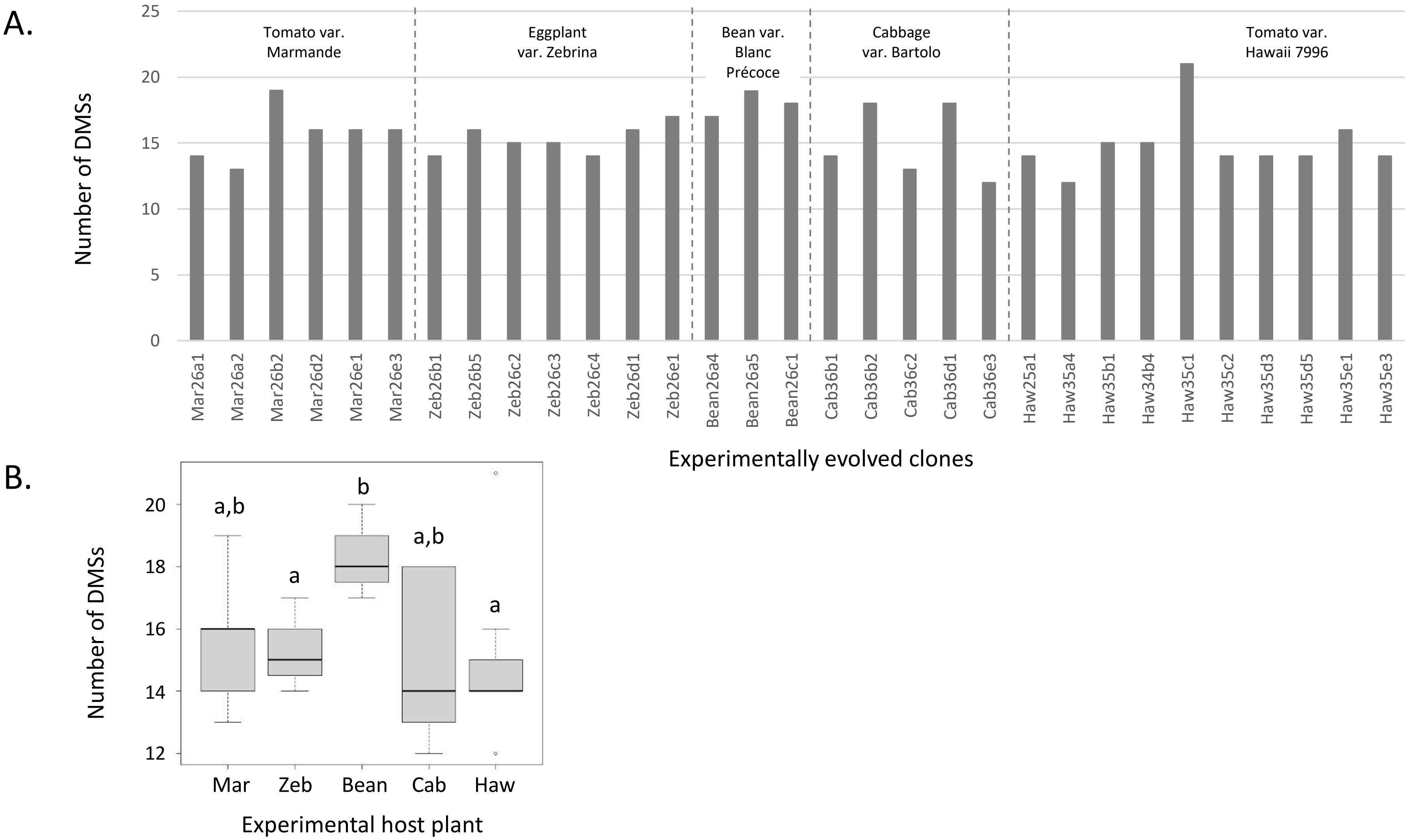


**Supplemental Figure 1** Effect of the experimental host on the number of differential methylated sites (DMSs) detected in the evolved clones according to SMRT-sequencing. A. Number of DMSs in each investigated evolved clone. B. Mean number of DMSs in evolved clones for each experimental host. Different letters above the boxplot indicate a significant difference (Wilcoxon test, *p*.value $< 0.05$). Mar: Tomato var. Marmande; Zeb : Eggplant var. Zebrina; Bean : Bean var. Blanc précoce; Cab : Cabbage var. Bartolo; Haw: Tomato var. Hawaii 7996.
